## Supplementary Information for "Generation of human iPSC-derived phrenic-like motor neurons to model respiratory motor neuron degeneration in ALS"

### Supplementary Figures

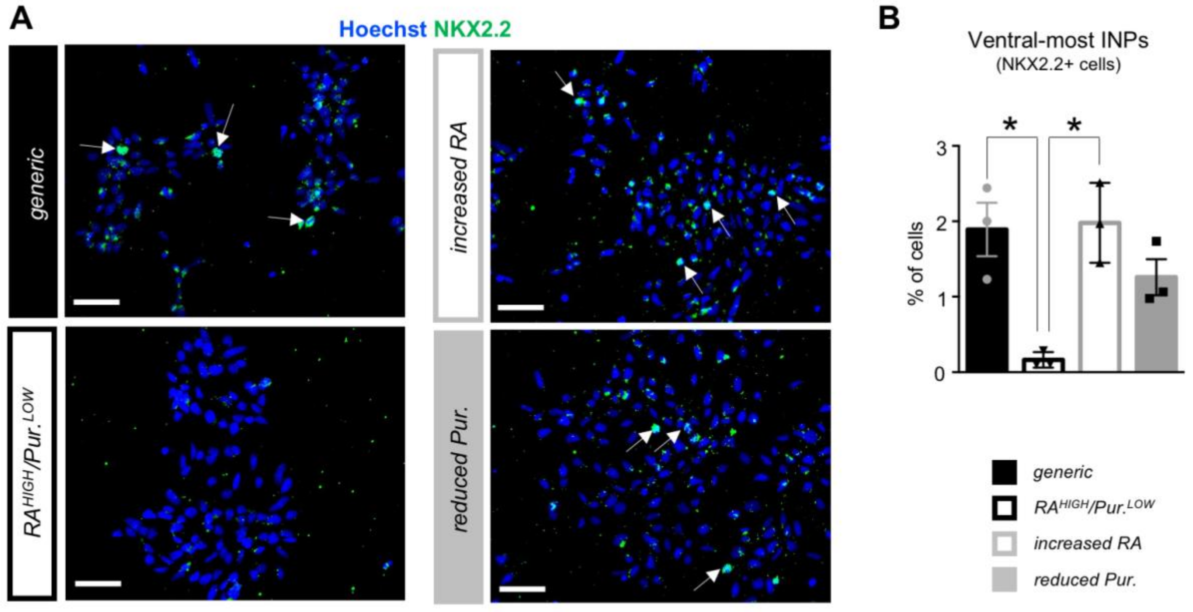


**Supplementary Figure 1. Dorsalization of the culture. (A)** Representative images of MNPC cultures stained with the anti-NKX2.2 antibody after 12 days of differentiation in each of the four experimental conditions defined in Figure 1A. **(B)** Quantification of the proportion of ventral-most interneuron progenitors identified as NKX2.2+ cells. Scale bars= 50 µm. Friedman non parametric test and Dunn's post-hoc multiple comparisons test; *=p<0.05; N= 3 cultures (with >500 cells in random fields for each culture).

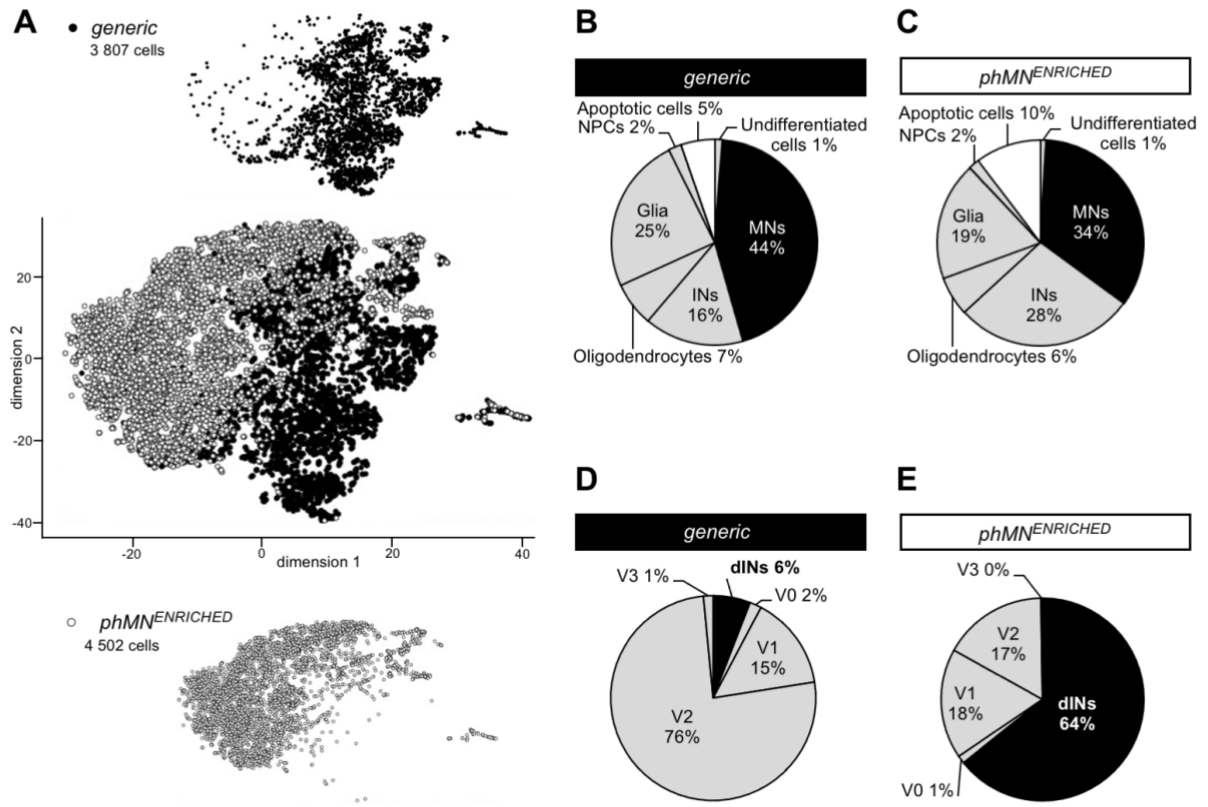


**Supplementary Figure 2. Heterogenous composition of human iPSC-derived MN culture defined by single-cell RNA sequencing. (A)** t-SNE plot of hiPSC derived MN cultures after 28 days of differentiation in the *generic* (black dots) or the *phMN*$^{ENRICHED}$ (white dots) condition. **(B-C)** Relative proportion of undifferentiated cells, NPCs, INs, MNs, astrocytic glial cells, oligodendrocytes, and apoptotic cells, identified after 28 days of differentiation in the *generic* condition **(B)** or with *phMN*$^{ENRICHED}$ **(C)**, based on the specific expression of the following genes: *NANOG* and *OCT4* for undifferentiated cells; *SOX1*, *SOX2*, and *MKI67* for NPCs; *PAX2*, *PAX3*, *LBX1*, *EVX1*, *EN1*, *CHX10*, *GATA3*, *SOX14*, *SIM1*, *TLX3* for INs; *NEUROG2*, *OLIG2*, *HB9*, *ISL1*, *ISL2*, *CHAT* for MNs; *S100B and SOX9* for astrocytic glial cells; *PDGFR*$\alpha$ and *GALC* for oligodendrocytes; *BCL2*, *BAX*, and *NGF* for apoptotic cells. For both conditions, the proportion of MNs is represented in black. **(D-E)** Proportion of each of the spinal interneuron (IN) subtypes, identified after 28 days of differentiation in the *generic* **(D)** or the *phMN*$^{ENRICHED}$ **(E)** condition, based on combinatorial gene expression: dorsal INs (dINs), *LBX1*$^{+}$/*PAX2*$^{+}$/*TLX3*$^{+}$; V0 INs, *EVX1*$^{+}$/*EN1*$^{-}$; V1 INs, *EVX1*$^{-}$/*EN1*$^{+}$; V2 INs, *CHX10*$^{+}$/*SOX14*$^{+}$/*GATA3*$^{+}$; V3 INs, *SIM1*$^{+}$/*NKX2.2*$^{+}$. For both conditions, the proportion of dINs is represented in black.

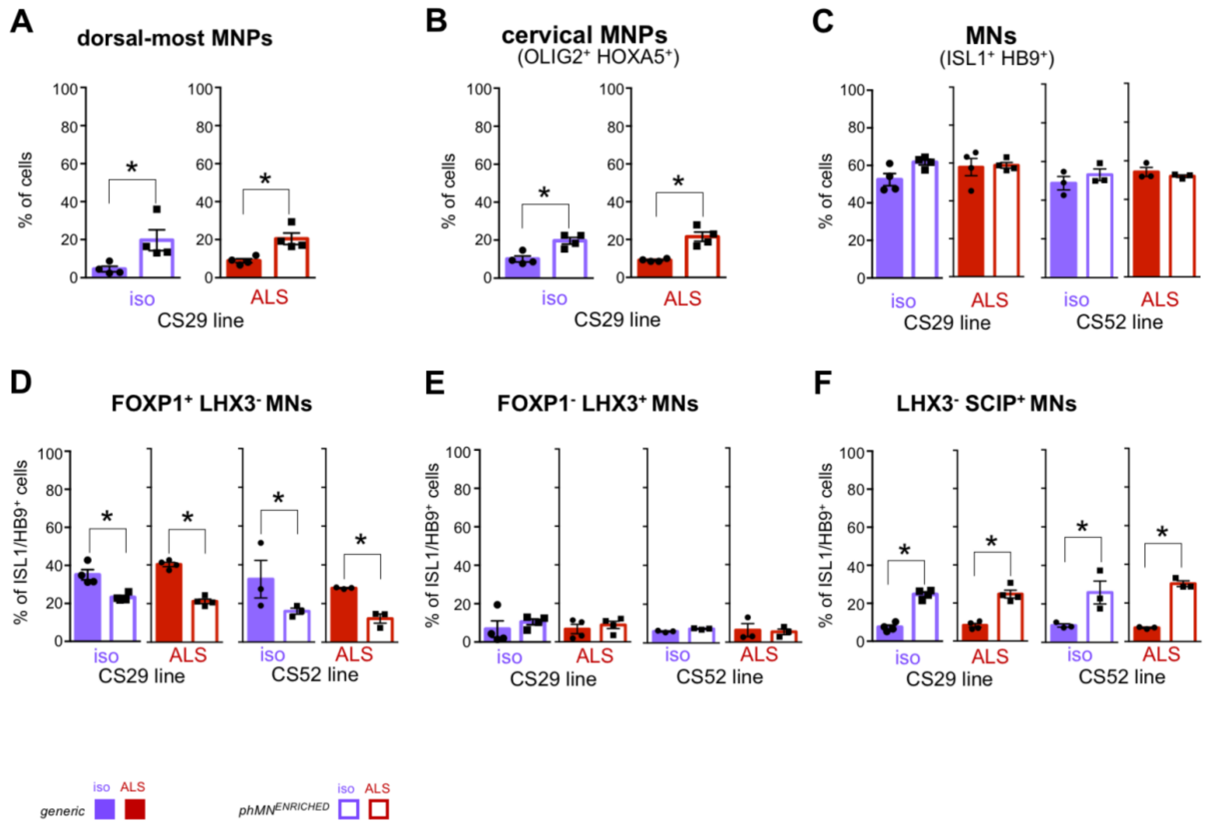


**Supplementary Figure 3. Enrichment in dorsal-most cervical-HMC MNPCs and phMNs derived from two C90rf72 iPSC lines. (A-B)** Quantification of the dorsal-most MNPCs (e.g., PAX6$^{high}$/TLE$^{low}$ OLIG2$^{+}$ cells, (**A**) and the cervical MNPCs (e.g., HOXA5+/OLIG2+ cells, (**B**) as percentages of the total number of cells after 12 days of differentiation in either the *generic* or the *phMN$^{ENRICHED}$* culture conditions established in previous figures. **(C)** Quantification of MNs (e.g., ISL1/HB9$^{+}$ cells) as percentages of the total number of cells after 25 days of differentiation. **(D-F)** Quantification of the LMC MNs identified as FOXP1$^{+}$/LHX3$^{-}$ **(D)**, MMC MNs identified as LHX3$^{+}$/FOXP1$^{-}$ **(E)**, and phMNs identified as LHX3$^{-}$/SCIP$^{+}$ **(F)** as percentages of the number of ISL1/HB9$^{+}$ cells. Scale bars= 50 µm. N= 3 cultures (with >500 cells in random fields for each culture). MMC= median motor column, LMC= lateral motor column, phMNs= phrenic MNs. Mann-Whitney non parametric test; *=$p<0.05$; N= 4 cultures for CS29 and N=3 cultures for CS52 (with >500 cells in random fields for each culture).

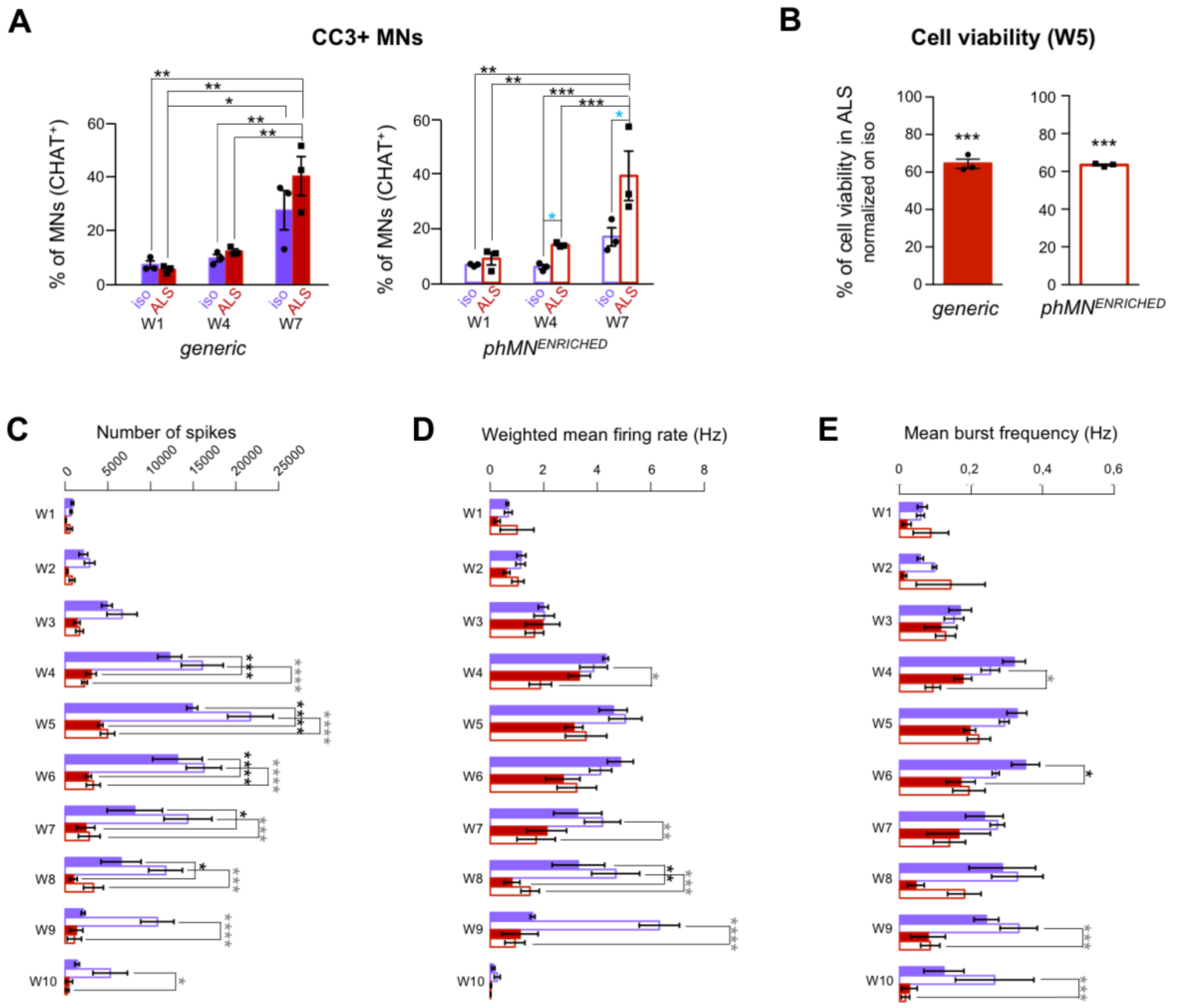


**Supplementary Figure 4. C9orf72 ALS MN death in 'generic' or 'enriched' cultures derived from the CS52 hiPSC line. (A)** Quantification of the $CC3^+$ MNs (e.g., $CC3^+/CHAT^+$ cells) as percentages of the total number of MNs (e.g., CHAT+ cells) in *generic* (left, filled bars) or *phMN*$^{ENRICHED}$ (right, hollow bars) cultures derived from the CS52 iPSC line. Two-way ANOVA and Tukey's post-hoc multiple comparisons test; *=p<0.05; **=p<0.005; ***=p<0.0005; N= 3 cultures per condition. Blue asterisks show statistical significance between Iso and ALS at the same time point. **(B)** Cell viability assay measured using CellTiter-GLO after 60 days of differentiation (i.e., five-weeks (W5) post-plating) in *generic* (left, filled bars) or *phMN*$^{ENRICHED}$ (right, hollow bars) cultures derived from the CS52 iPSC line. Cell viability for ALS cells was normalized on each respective corresponding isogenic control. Wilcoxon signed rank test; ***=p<0.0001; N= 3 cultures per condition. **(C-E)** Histograms showing the average number of spikes (**C**)**,** weighted mean firing rate (Hz) (**D**), and mean burst frequency (Hz) (**E**) over time, from one-week to 10-weeks post-plating of isogenic *generic*, isogenic *phMN*$^{ENRICHED}$, ALS *generic*, and ALS *phMN*$^{ENRICHED}$ from the CS52 hiPSC line. Two-way repeated measure ANOVA and Tukey's post-hoc multiple comparisons test; *=p<0.05; **=p<0.005; ***=p<0.0005; ****=p<0.0001; N= 3 cultures per condition. Black asterisks = statistical significance for isogenic vs ALS *generic* cultures; Grey asterisks = statistical significance for isogenic vs ALS *phMN*$^{ENRICHED}$ cultures.

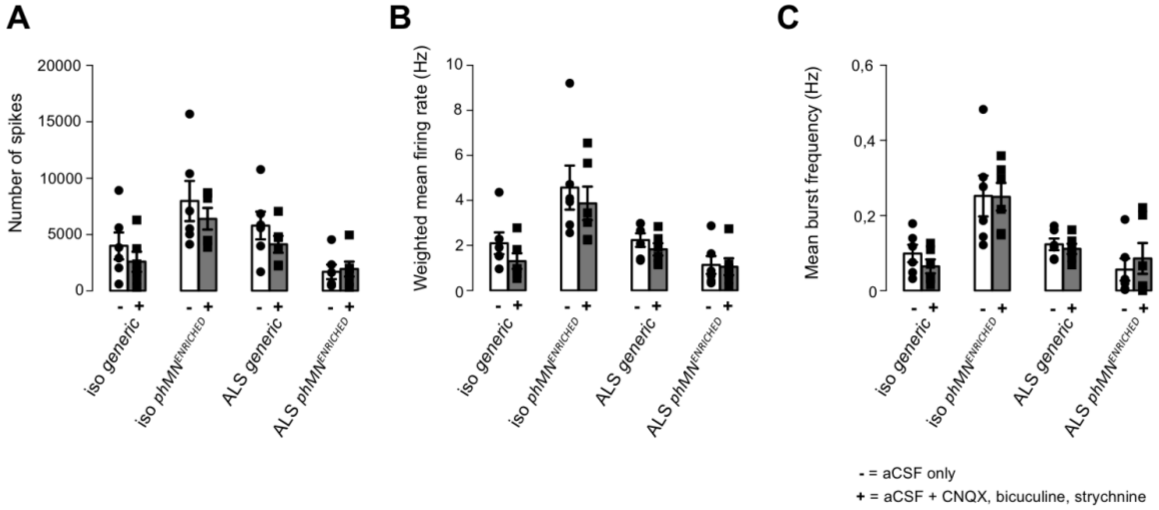


**Supplementary Figure 5.** Histograms showing the average number of spikes (**A**)**,** weighted mean firing rate (Hz) (**B**), and mean burst frequency (Hz) (**C**) at 5-weeks (D60) post-plating for each of the four culture conditions derived from the CS29 line in the absence (white bars; '-') or presence (grey bars; '+') of drugs suppressing all synaptic currents (CNQX, bicuculine and strychnine). Mann-Whitney non parametric test comparing data in the absence or presence of drugs; no statistically significant difference; N=6 cultures per condition.
